# Analysis of a meningococcal meningitis outbreak in Niger - potential effectiveness of reactive prophylaxis

**DOI:** 10.1101/496299

**Authors:** Matt D.T. Hitchings, Matthew E. Coldiron, Rebecca F. Grais, Marc Lipsitch

## Abstract

**Background:** Seasonal epidemics of bacterial meningitis in the African Meningitis Belt carry a high burden of disease and mortality. Reactive mass vaccination is used as a control measure during epidemics, but the time taken to gain immunity from the vaccine reduces the flexibility and effectiveness of these campaigns. Highly targeted reactive antibiotic prophylaxis could be used to supplement reactive mass vaccination and further reduce the incidence of meningitis, and the potential effectiveness and efficiency of these strategies should be explored.

**Methods and Findings:** Data from an outbreak of meningococcal meningitis in Niger, caused primarily by *Neisseria meningitidis* serogroup C, is used to estimate clustering of meningitis cases at the household and village level. In addition, reactive antibiotic prophylaxis and reactive vaccination strategies are simulated to estimate their potential effectiveness and efficiency, with a focus on the threshold and spatial unit used to declare an epidemic and initiate the intervention.

There is village-level clustering of meningitis cases after an epidemic has been declared in a health area. Meningitis risk among household contacts of a meningitis case is no higher than among members of the same village. Village-wide antibiotic prophylaxis can target secondary cases in villages: across of range of parameters pertaining to how the intervention is performed, up to 200/ 672 cases during the season are potentially preventable. On the other hand, household prophylaxis targets very few cases. In general, the village-wide strategy is not very sensitive to the method used to declare an epidemic. Finally, village-wide antibiotic prophylaxis is potentially more efficient than mass vaccination of all individuals at the beginning of the season, and than the equivalent reactive vaccination strategy.

**Conclusions:** Village-wide antibiotic prophylaxis should be considered and tested further as a response against outbreaks of meningococcal meningitis in the Meningitis Belt, as a supplement to reactive mass vaccination.

## Author summary

Until a low-cost polyvalent conjugate meningococcal vaccine becomes available in the African Meningitis Belt, reactive strategies to control meningitis epidemics should be considered and tested, and refined in order to maximise effectiveness. A recent cluster-randomised trial conducted in Niger showed promising evidence for the effectiveness of a village-wide reactive antibiotic prophylaxis intervention. We used data from a meningitis outbreak in Niger to explore the potential effectiveness and efficiency of this and other strategies when deployed on a wider scale, allowing us to compare different strategies without recourse to further randomised trials. This study provided further evidence that village-wide antibiotic prophylaxis targets secondary cases in villages, and showed that the intervention remains effective whether it is initiated early in the season (targeting more cases during the season) or later (when clustering of cases by village is strongest). For this outbreak, reactive village-wide antibiotic prophylaxis would have been more potentially efficient than mass vaccination at the beginning of the season, implying that targeted prophylaxis could supplement reactive mass vaccination. Many authors have developed models for vaccination strategies to reduce the burden of meningitis in sub-Saharan Africa; our results add to this literature by considering antibiotic prophylaxis as a supplementary intervention.

## Introduction

Epidemics of bacterial meningitis occur seasonally in the “Meningitis Belt” of sub-Saharan Africa, and are most commonly due to *Neisseria meningitidis*(1, 2). In Niger and throughout the Meningitis Belt, spatial clustering of cases(3, 4) can be partly but not fully explained by variations in climatic factors, suggesting the role of the environment and transmission in driving epidemics(5).

Individuals in close contact with meningitis cases are at higher risk for carriage of *N. meningitidis* and invasive disease, among epidemic and non-epidemic settings(6, 7). Household contacts of meningococcal meningitis cases are at higher risk of meningococcal meningitis than the general population, and the risk ratio has been reported to be as high as 1,000(8, 9). In high-resource settings, the effectiveness of household chemoprophylaxis has been estimated to reduce the risk of meningitis by 84%(10).

Antibiotic prophylaxis of household members of meningococcal meningitis cases is recommended by the World Health Organisation (WHO) in sub-Saharan Africa outside of an epidemic only(11). This is because meningitis burden and carriage prevalence are much higher during epidemics(12), so household chemoprophylaxis would be labor-intensive and could have minimal impact on overall carriage.

The MenAfriVac conjugate vaccine provides long-lasting protection against carriage, leading to vast reductions in the burden of meningitis due to *N. meningitidis* serogroup A (NmA) since its introduction. However, polysaccharide vaccines available in the Meningitis Belt against other serogroups provide only short-lived protection against disease. Until a low-cost conjugate vaccine targeting these serogroups becomes widely available, reactive mass vaccination campaigns using polysaccharide vaccines can be conducted during epidemics. However, they are difficult to organize and implement in a timely fashion, and thus their impact in reducing cases can be limited(13). Targeted prophylactic interventions at a smaller spatial scale could lead to further reduction in cases during epidemics. A recent cluster-randomized trial in Niger during an outbreak of meningitis caused by NmC found promising evidence for the effectiveness of village-wide prophylaxis with single-dose ciprofloxacin at reducing the incidence of meningococcal meningitis at the community level. Overall incidence was not reduced when prophylaxis was limited to household members of cases(14).

Several papers have examined the effect of different intervention thresholds on effectiveness of interventions for seasonal meningitis outbreaks(15–18). These studies have focused on reactive vaccination, which typically has a lag time of weeks between crossing the epidemic threshold to implementation. Antibiotic prophylaxis can be performed more quickly than vaccination and without the need for a cold chain, and antibiotics can be stockpiled more easily and cheaply. In addition, an individual receiving prophylaxis would receive protection immediately, and although this protection is unlikely to be as long-lasting, evidence suggests that ciprofloxacin is effective at clearing carriage up to two weeks after treatment(19).

To build on the promising results of the recent trial, it is important to understand the potential for reactive antibiotic prophylaxis to be used on a wide scale to supplement reactive mass vaccination and before a polyvalent conjugate vaccine is available. To this end, data from a single epidemic in the Dosso Region of Niger is used to describe clustering of cases at the household and village level, and estimate the potential effectiveness of several prophylaxis strategies.

## Methods

### Data Sources

Passive surveillance data from the 2015 meningitis season was collected (Fig 1). This secondary analysis was classified as exempt by the Harvard T.H. Chan School of Public Health IRB (ref: IRB17-0974), and all data analysed were anonymised. This season saw a large and unexpected outbreak of *N. meningitidis* serogroup C in Niger with 8,500 suspected cases reported(20). The peak was between 4-10 May, and the majority of cases were in Niamey in the southwest, followed by the Dosso Region, comprising 8 departments. This database was augmented by household follow-up visits to notified cases in the Dogondoutchi and Tibiri departments in September 2015, by which cases were linked by household, and household size was collected(21). Population and coordinates of the villages were sourced from the 2012 census and OpenStreetMap. The study area is made up of four departments (Dogondoutchi, Tibiri, Gaya, and Dioundiou) each of which is made up of communes (18 in total). In addition, health areas (*aires de santé*) are defined as the area served by a particular health centre. There are 38 health areas in the study area, with populations ranging from 8,000 to 56,000.

**Fig 1.**
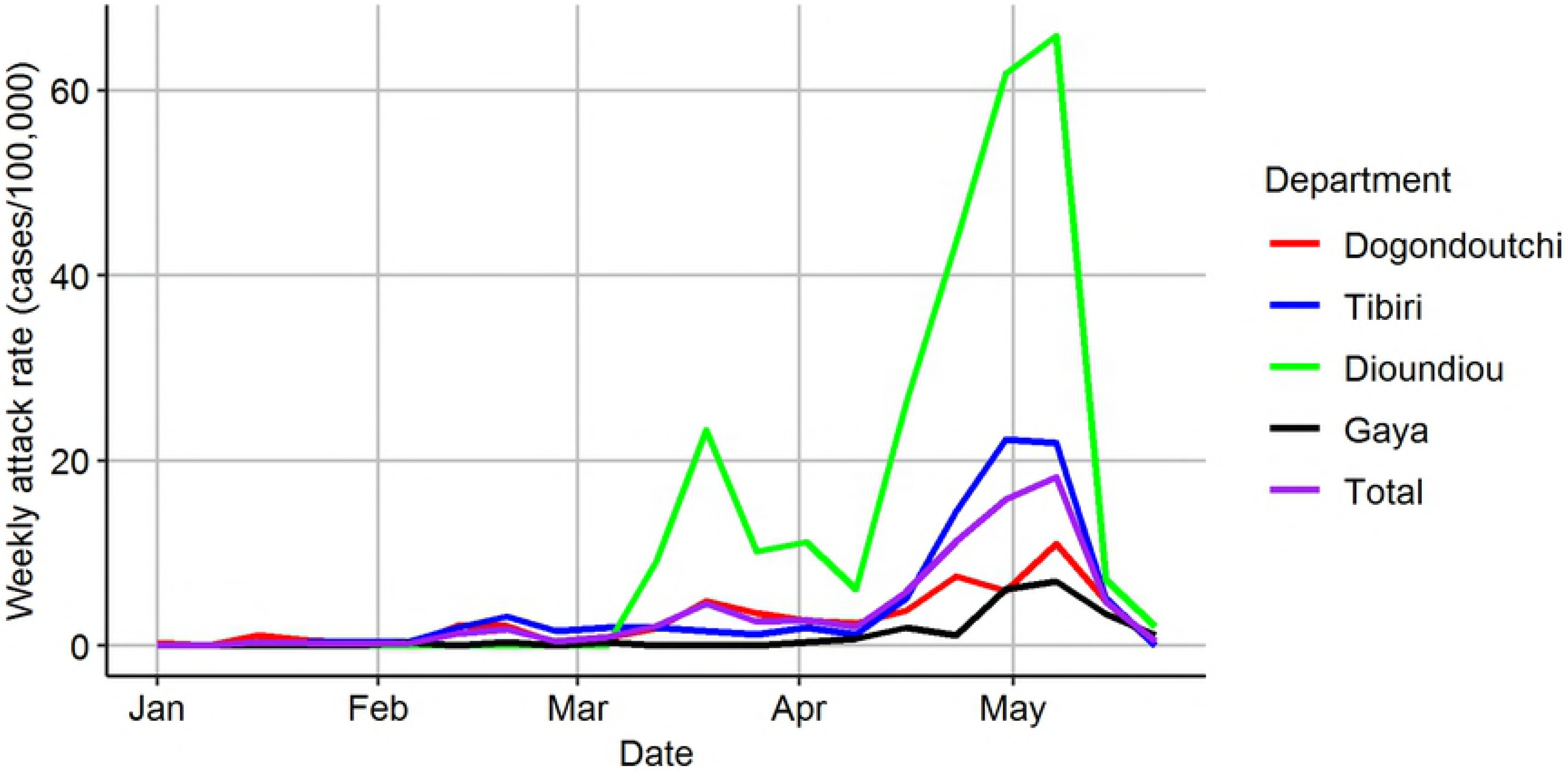
Epidemic curve in study area. Weekly attack rate in Dogondoutchi (red), Tibiri (blue), Dioundiou (green), Gaya (black), and in the whole study area (purple).

The data base contains patient-level information on 752 suspected cases in the Dogondoutchi, Tibiri, Gaya and Dioundiou departments between January 2 and May 23. After excluding cases whose origin was in Nigeria, 348/429 cases in Dogondoutchi and Tibiri (81%) were reached for the household survey. The census data base contained data on 2,588 villages, with 310 villages appearing in the case data. The population and coordinates of 246 out of those 310 villages were obtained, representing 689 cases (92%). Of these villages, 26 were neighborhoods of the larger cities of Dogondoutchi (total population 27,427) and Gaya (total population 44,809). 495 (66%) of cases had cerebrospinal fluid samples tested, of which 291 (59%) were confirmed *N. meningitidis* (serogroup C, W, or unspecified), 17 (3%) were confirmed *S. pneumoniae,* and 187 (38%) tested negative for the presence of these two bacteria.

Table 1 shows the variability in commune size, number of suspected cases and whether and when the epidemic threshold was crossed.

**Table 1.**
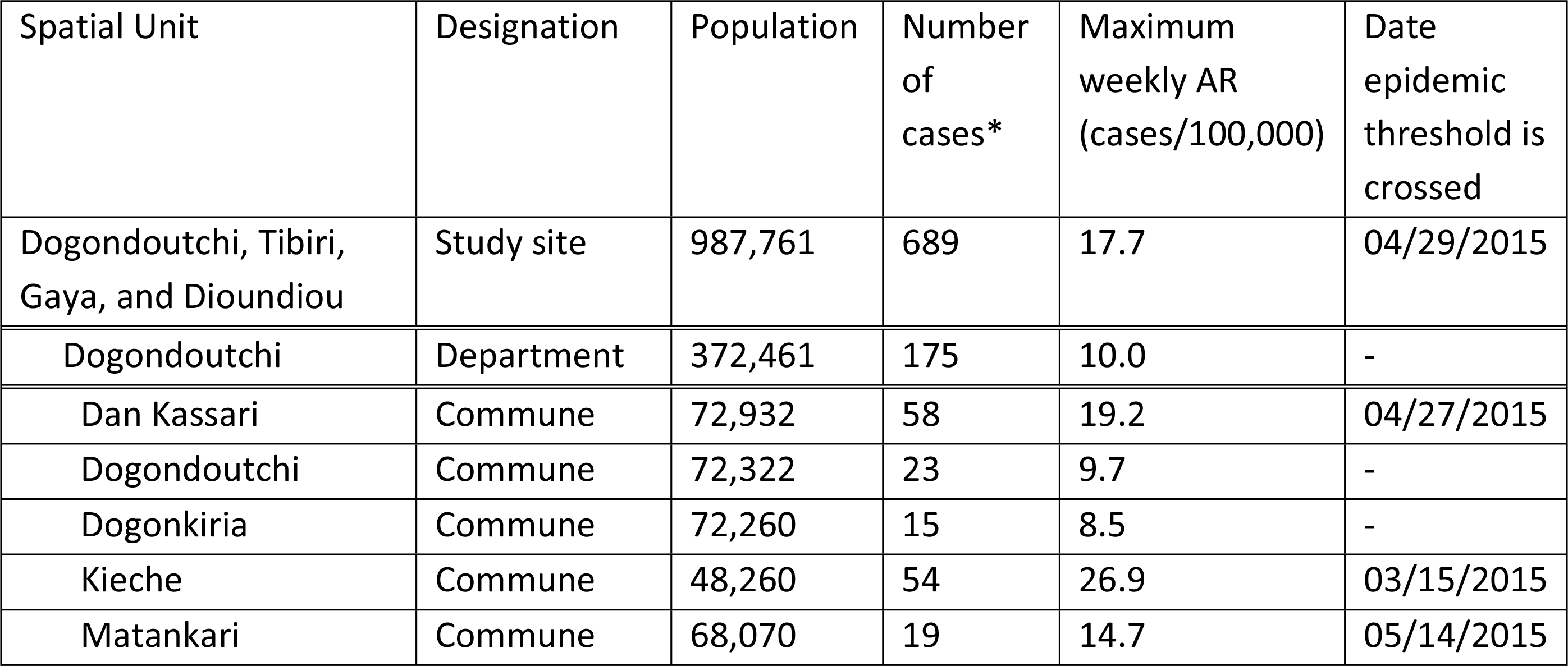
Number of cases and population across study area.

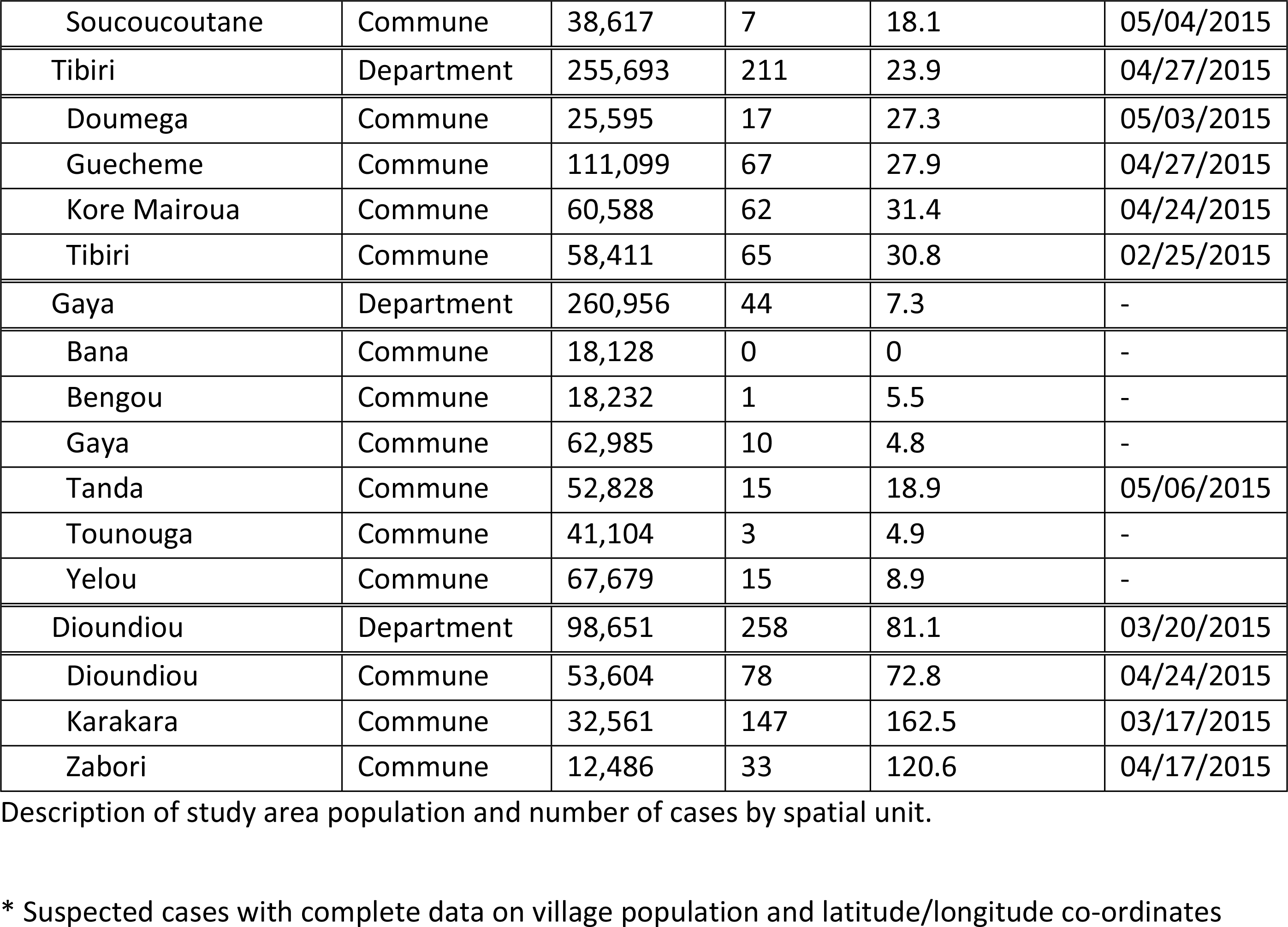

### Definitions

A *N. meningitidis* epidemic is defined by whether the weekly attack rate (cases/100,000) has reached a certain threshold(8). The current epidemic threshold used by the WHO is 10 cases/100,000 for any population greater than 30,000, or 5 cases in a week for any population under 30,000. We apply thresholds of 3, 5, 7, and 10 cases/100,000 to three spatial units: health area, commune, and health district, to define whether a region is in an epidemic or not.

We are interested in clustering of cases at two spatial units: the household and the village. We define a “contact” of a case as a member of the spatial unit of interest, specifically a household member or resident of the same village. Specifically, an individual is defined as a “contact” of a case if a suspected case has previously occurred in their spatial unit.

### Clustering measures

Clustering at the household and village level is described by calculating two metrics:

- the relative risk of meningococcal meningitis for a contact of a suspected meningococcal meningitis case compared to a non-contact (defined as the “household relative risk (RR)” or “village relative risk (RR)”);
- and the proportion of cases that are contacts of a suspected case.

The household RR is presented unadjusted and adjusted for the village-level cumulative incidence. Villages with higher attack rates are more likely to have households with multiple cases by chance, and therefore the unadjusted household RR, while useful from a policy standpoint as it identifies high-risk individuals in the population, is biased upwards in describing the relative risk that might be causally due to having a household contact. Similarly, the village RR is adjusted for the commune-level cumulative incidence. The question of whether the pattern of clustering is different outside of an epidemic compared to during an epidemic is addressed by defining such periods and comparing the metrics by outside/during epidemic status.

Household RR is estimated using Poisson regression with rate of meningococcal meningitis as the outcome, and household contact as the exposure of interest. We controlled for the cumulative incidence of meningococcal meningitis in the village across the follow-up period by including log(cumulative incidence) as a variable in the regression model. To compare the household RR in the non-epidemic and epidemic period, we categorized all cases as “epidemic” or “non-epidemic” according to the current WHO epidemic threshold applied at the health area level. We report the household RR in the non-epidemic and epidemic period separately, as well as the relative household RR and confidence interval. Village RR is calculated in a similar way.

The proportion of cases that are contacts of a previously-notified case was calculated, and a confidence interval was estimated using log-binomial regression among cases only, with “having a contact” as the outcome. To assess whether the proportion changes between the non-epidemic and epidemic period, we include it as a variable in the model as described above.

### Reactive prophylaxis intervention

We simulated a variety of prophylaxis strategies on the data, restricted to rural villages only (i.e. those that were not neighborhoods in the cities of Dogondoutchi or Gaya).

We simulate the reactive prophylaxis strategy as follows (see Fig 2). The entire study area starts in the “pre-epidemic” state, in which surveillance for meningococcal meningitis cases is performed at the level of the *surveillance unit* (health area, commune, or department). When the attack rate has reached a given threshold in a surveillance unit, an epidemic is declared in that unit (as in the middle region in Fig 2). From this day onwards, the unit enters the “epidemic” state, in which villages in the unit are followed for the incidence of cases. When a triggering case occurs in a village (as in the second village in Fig 2), the village enters the “contact prophylaxis” state, in which all contacts of the triggering case are identified and provided prophylaxis. The contacts are defined either as household members, village members, or all members of villages within a certain radius of the triggering case’s village.

**Fig 2.**
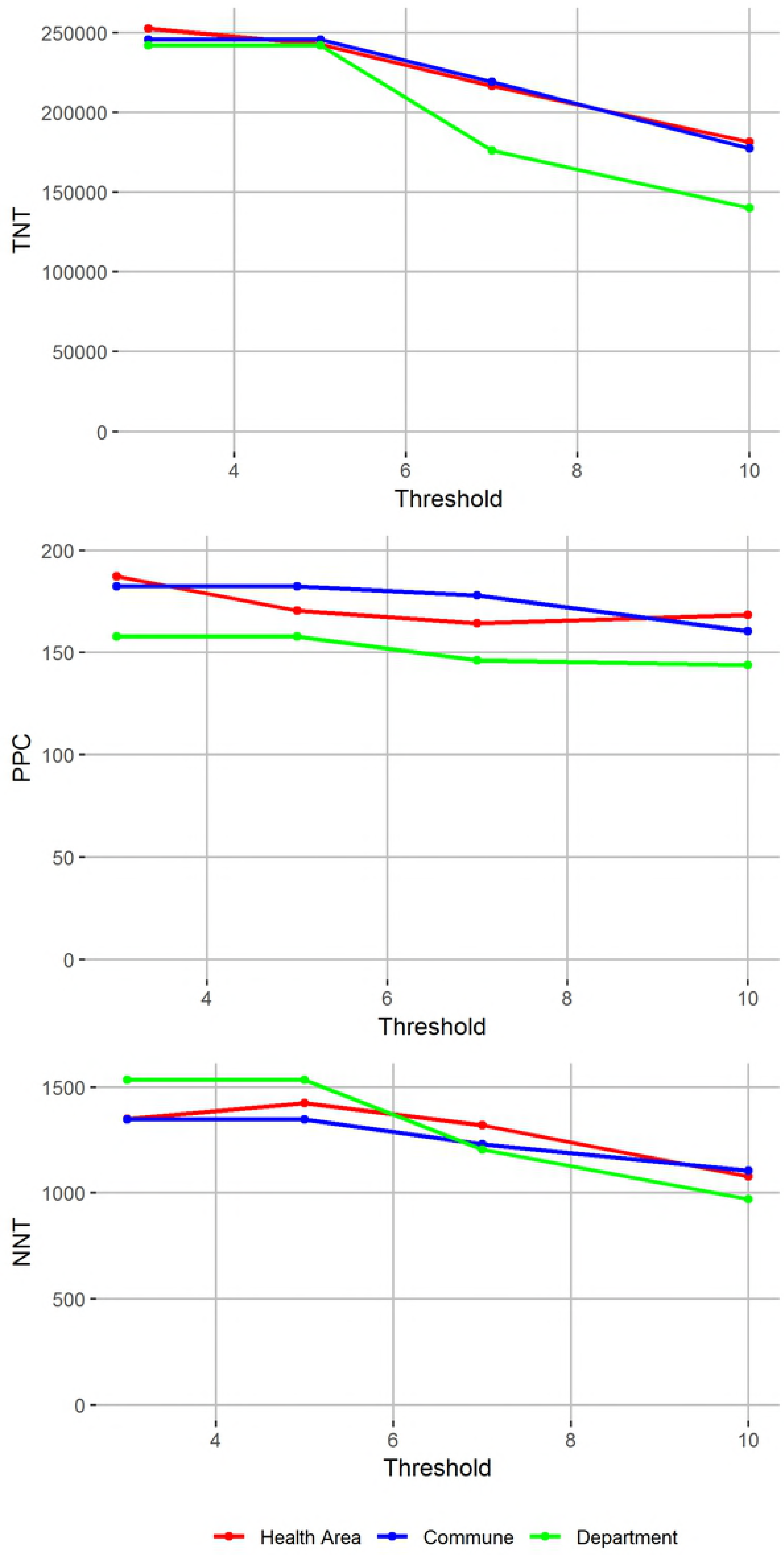
Schematic of the reactive prophylaxis protocol. Description of the reactive prophylaxis protocol, in the pre-epidemic, epidemic, and contact prophylaxis stages.

The number of doses needed for each contact prophylaxis is calculated using population data. The number of potentially prevented cases (PPC) from each round is defined as the number of cases that occur within a given time window after antibiotic distribution, and the total PPC is the sum of PPCs from all contact prophylaxis rounds conducted during the intervention. In performing this analysis, we focus on the direct effect of prophylaxis only, and make no assumptions about indirect effects caused by clearing of carriage from targeted contacts. The total number treated (TNT) is the total number of doses administered. The number needed to treat (NNT) per potentially prevented case is calculated as NNT=TNT/PPC. Once a village is given a round of prophylaxis, cases that occur in that village during the presumed time window of effectiveness do not trigger new rounds of prophylaxis, although cases that occur after the end of the window can trigger further rounds (which is relevant for the radial strategies, or if villages are repeatedly treated).

### Reactive vaccination

Finally, we simulate a reactive vaccination strategy as a comparison for the chemoprophylaxis strategies. As well as simulating the above strategies, we calculate the PPC and number needed to vaccinate (NNV) for a strategy in which mass vaccination of the entire study area is conducted on the day the first case occurs in the season. While this strategy is unrealistic, it represents the best possible strategy in terms of PPC and serves as a basis of comparison for the other interventions.

Table 2 shows parameters in the model, meanings, and values considered for simulations. We consider all suspected cases, excluding only cases that tested positive for *S. pneumoniae.* In effect, we assume that all cases that tested negative for *N. meningitidis* are in fact false negatives. We perform a sensitivity analysis in which we exclude cases that test negative, and assume that the proportion of untested cases that are positive for *N. meningitidis* is equal to the proportion of tested cases that are positive. Given the uncertainty around the serial interval for *N. meningitidis* and other mechanisms of protection granted by prophylaxis, we assume a range of time windows during which prophylaxis can prevent cases. The evidence for the effectiveness of prophylaxis is strongest for cases occurring in the two weeks following index case identification(14, 19). We assume that during the course of the season, no individual can be treated more than once, although we relax this assumption in a sensitivity analysis. In addition, we consider strategies in which only villages below a certain population size are targeted. We make no assumptions about the efficacy of prophylaxis, reporting only the cases that could be targeted within a given time window.

**Table 2.**
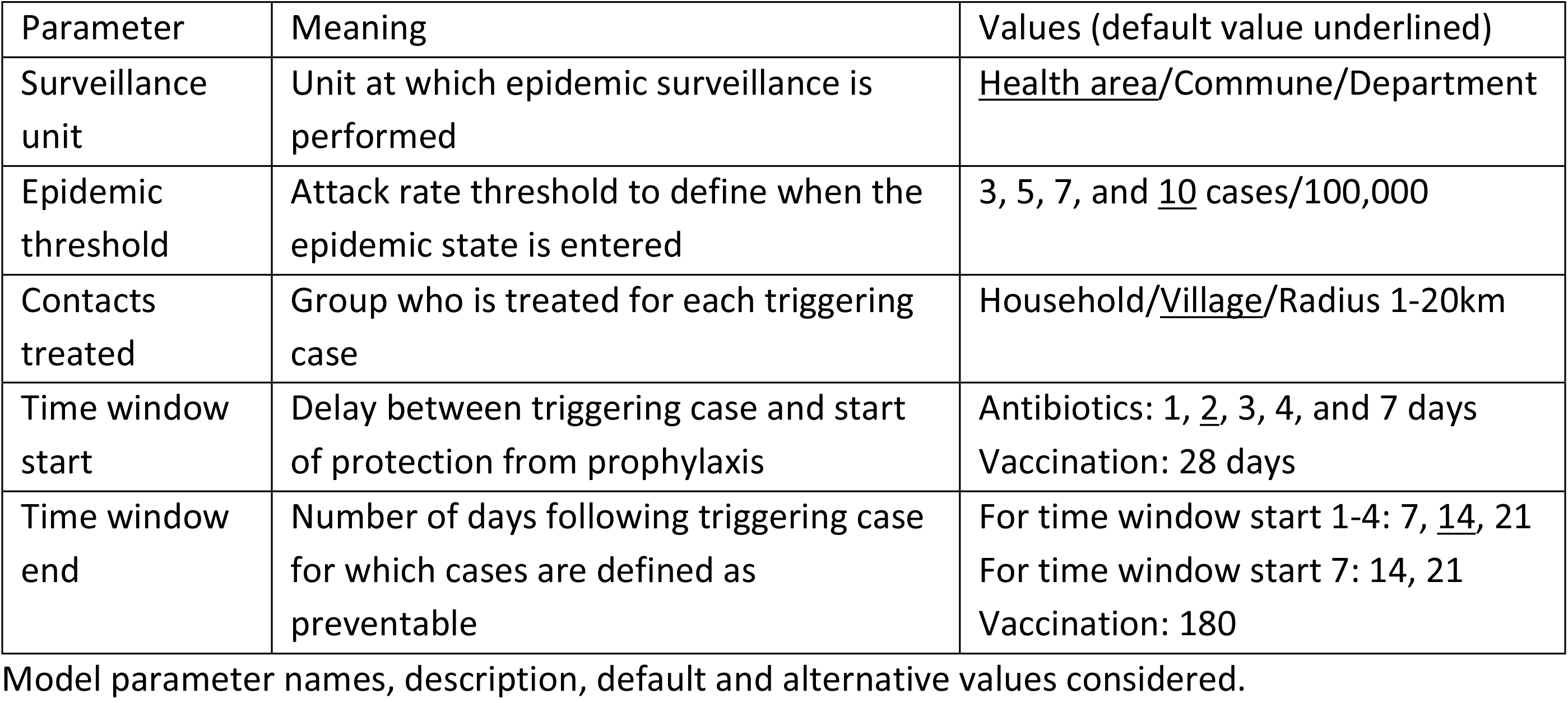
Parameters, meanings, and values considered.

| Parameter | Meaning | Values (default value underlined) |
| --- | --- | --- |
| Surveillance unit | Unit at which epidemic surveillance is performed | <u>Health area</u>/Commune/Department |
| Epidemic threshold | Attack rate threshold to define when the epidemic state is entered | 3, 5, 7, and <u>10</u> cases/100,000 |
| Contacts treated | Group who is treated for each triggering case | Household/<u>Village</u>/Radius 1-20km |
| Time window start | Delay between triggering case and start of protection from prophylaxis | Antibiotics: 1, 2, 3, 4, and 7 days<br>Vaccination: 28 days |
| Time window end | Number of days following triggering case for which cases are defined as preventable | For time window start 1-4: 7, 14, 21<br>For time window start 7: 14, 21<br>Vaccination: 180 |

Model parameter names, description, default and alternative values considered.

## Results

### Clustering

Clustering metrics at the village and household level are shown in Table 3. Household metrics were calculated using data only from those households that were reached for follow-up visits. The household secondary attack rate is nearly four times greater than the attack rate among individuals not exposed to a household contact rate. However, there is no elevated meningococcal meningitis risk to household members of a meningococcal meningitis case compared to other members of the same village. At the village level, members of a village with a meningococcal meningitis case have significantly elevated risk of meningococcal meningitis compared with other members of the same commune, and over 60% of cases occur in a village that has had a previous case.

**Table 3.** Household and village clustering metrics.

| Metric | Household | Village |
| --- | --- | --- |
| Relative risk | 3.91 (2.27, 6.24) | 3.12 (2.67, 3.64) |
| Relative risk (adjusted) | 0.93 (0.53, 1.52) | 2.09 (1.78, 2.46) |
| % cases that had a past contact | 5.0% (3.0%, 7.8%) | 62.1% (58.4%, 65.7%) |

Relative risk, adjusted relative risk, and proportion of secondary cases, estimated at the household and village level.

The point estimate of household relative risk is lower in the epidemic period than in the nonepidemic period, but the confidence intervals are wide and the difference is not significant (relative risk ratio 0.69, 95% CI(0.25, 2.06)), although there is a lack of power as only 16 secondary cases were included in the analysis. There is evidence for clustering by village is only during an epidemic: village RR is 4.80, 95% CI(3.92, 5.93) during an epidemic compared to 1.01, 95% CI(0.76, 1.33) in the non-epidemic period (see Table S1).

### Household prophylaxis

The household prophylaxis strategy, under baseline parameter values, would have prevented six cases, hampered by the fact that only 4% of cases could possibly be targeted by a household-based intervention. On the other hand, a village-wide prophylaxis strategy would have targeted 178 eventual cases under baseline parameter values. Even though the household strategy prevents a small number of cases, it is much more efficient than the village strategy, with an NNT of 259.5 compared to 1,020.3 per PPC.

### The effect of thresholds on village-wide prophylaxis

The combination of threshold for intervention and spatial unit at which the threshold is applied changes the number of cases targeted and efficiency of the village-prophylaxis strategy by determining on which day during the season each village receives its round of prophylaxis, and whether it receives any prophylaxis. Fig 3 shows the TNT, NNT and PPC for various combinations of threshold and intervention unit.

**Fig 3.**
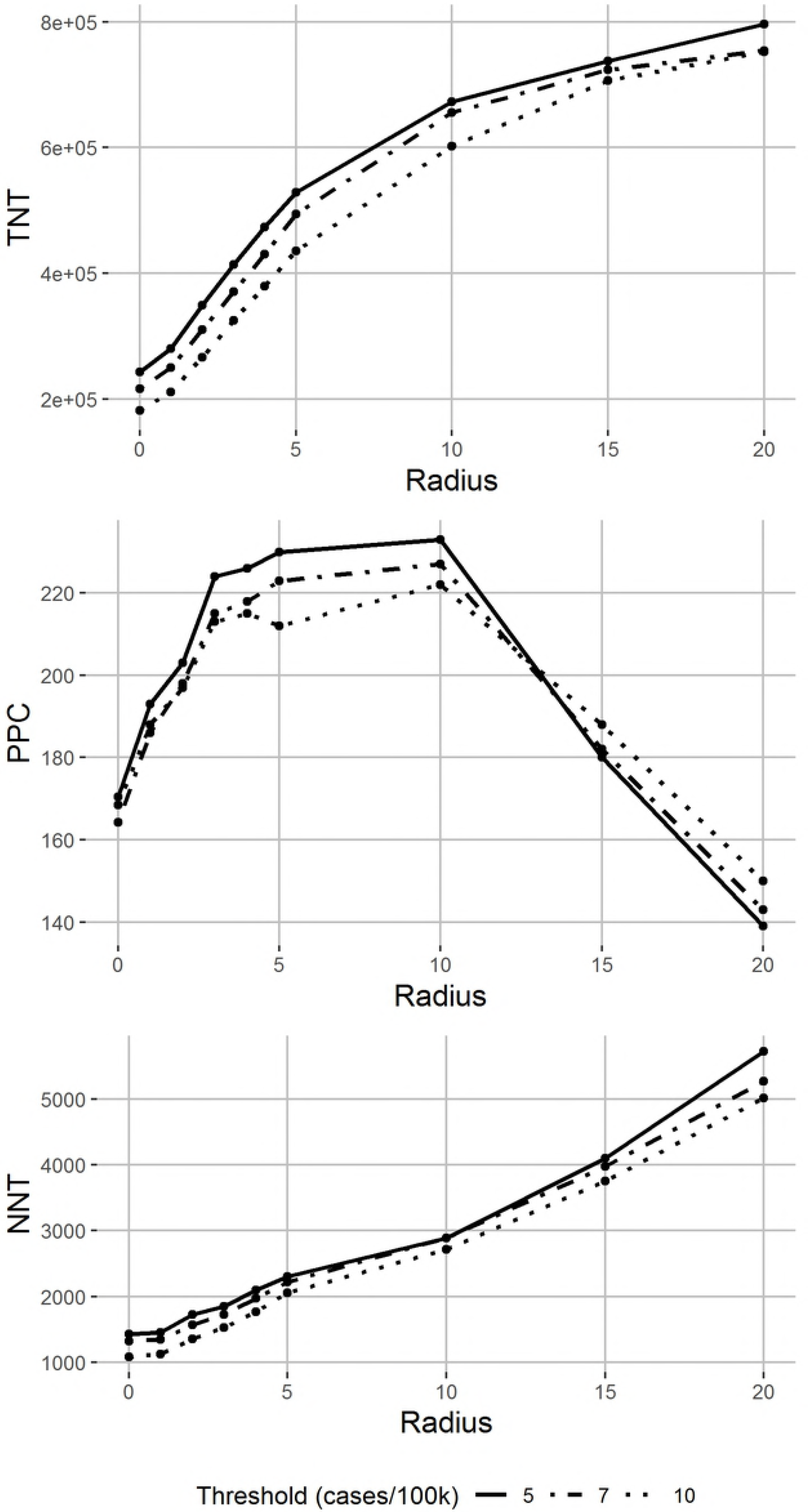
Potential effectiveness and efficiency of village-prophylaxis strategies by epidemic threshold definition. Total number treated, potentially prevented cases (PPC) and number needed to treat per PPC from applying a village-prophylaxis strategy, varying the threshold for intervention, with surveillance at different spatial units (colors).

As the threshold increases the PPC decreases because higher thresholds miss the opportunity to prevent clustered cases before the threshold is passed or in districts that never reach the threshold, while the TNT decreases because the intervention starts later in the season and some regions never pass the higher thresholds. On the other hand, the clustering is stronger later in the season, meaning that contacts of a case are at higher risk of meningococcal meningitis compared to non-contacts later in the season compared to earlier in the season. Therefore, NNT also decreases with threshold (Fig 3).

There are small differences between NNT and PPC across the three surveillance units. When surveillance is performed at the department level, interventions are initiated later in the season when clustering is strongest, so although NNT is lowest when surveillance is performed at the department level using a 10 cases/100,000 threshold, this strategy also prevents fewer cases.

### Radial prophylaxis strategies

Given that spatio-temporal clustering of cases has been shown in previous outbreaks, a prophylaxis strategy targeting multiple villages might be expected to potentially prevent more cases. However, if each village can only be targeted once in the season, a large radius might get “ahead” of the clustering and target villages too early to prevent cases. Whether this happens is determined by a combination of the spatial unit at which the threshold is monitored (health area, commune, or department), the radius of intervention, and the number of days prophylaxis can be expected to protect cases.

This logic is borne out in Fig 4, in which TNT, NNT and PPC are shown by radius of the treatment unit, for thresholds of 5, 7, and 10 cases/100,000 applied at the health area level. A radius of 10km around the triggering case increases the PPC relative to the village approach. A higher radius targets villages that experience cases after the prophylaxis window, and the PPC decreases as the radius increases from 10 to 20km. In general, increasing radius leads to increasing TNT, as more villages with no cases are targeted. NNT also increases with radius, as the population-level attack rate is low and only 310 out of 2,588 villages (12%) experience any cases. The above pattern is similar when the threshold is monitored at the commune and department level.

**Fig 4.**
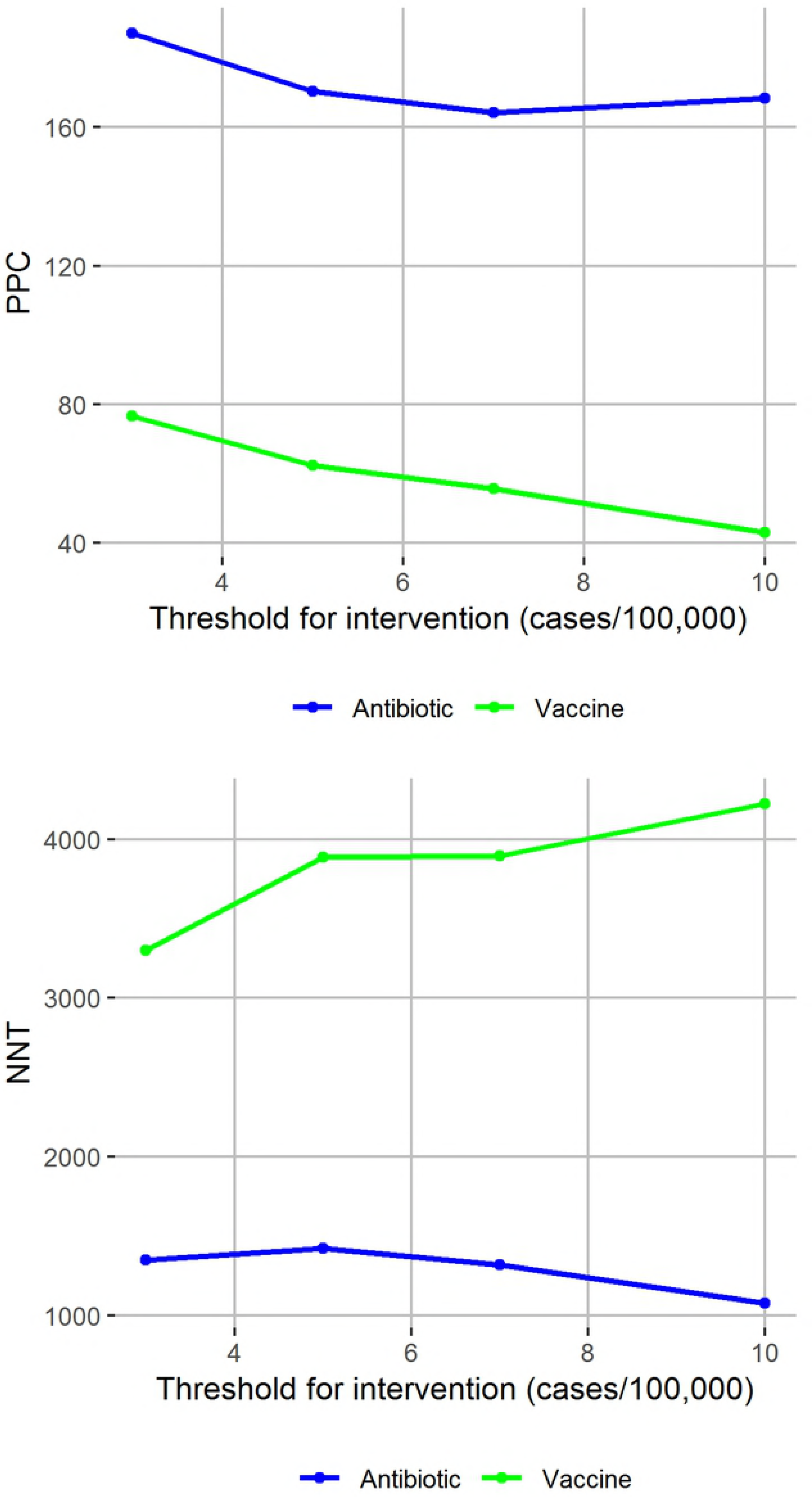
Potential effectiveness and efficiency of prophylaxis strategies by radius of prophylaxis. Total number treated, potentially prevented cases (PPC) and number needed to treat per PPC by radius of prophylaxis, varying the health area-level threshold for intervention start (line type).

### Comparison of reactive vaccination and reactive village prophylaxis

The most effective possible strategy, mass vaccination of the study population upon notification of the first case in the season, would have targeted 645 PPC, with NNV of 1531.4 vaccines per PPC. Other more targeted reactive vaccination strategies would have been much less effective at targeting cases due to the lag between case notification and implementation of the vaccination strategy (Fig 5), and the speed of an epidemic within a single village.

**Fig 5.**
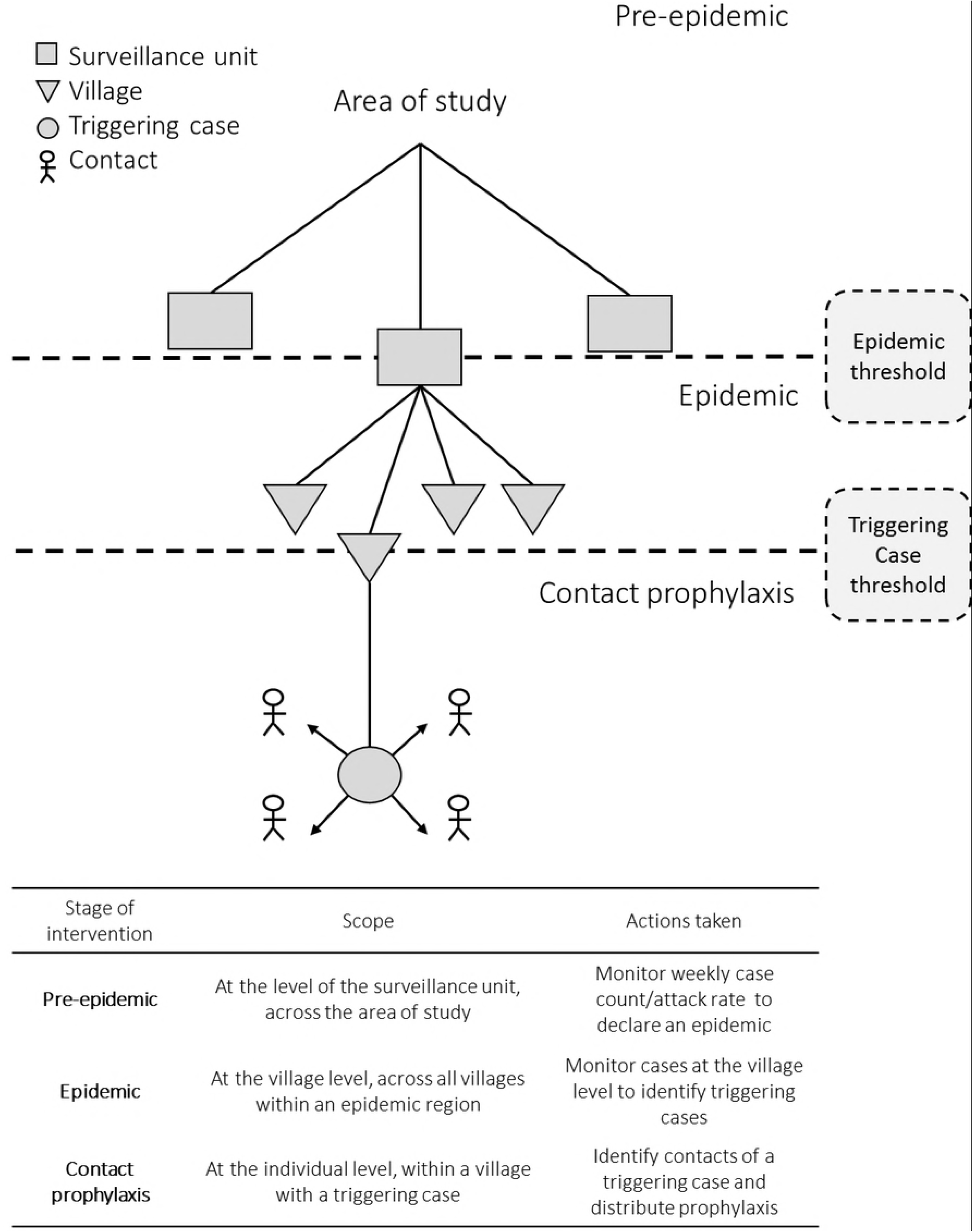
Potentially effectiveness and efficiency of village-antibiotic and village-vaccination prophylaxis strategies. Potentially prevented cases (PPC) and number needed to treat/vaccinate per PPC from applying a village-prophylaxis antibiotic (blue) and vaccination (green) strategy, varying the threshold for intervention at the health area level.

Under baseline parameter values, a village-wide reactive antibiotic prophylaxis strategy targets between 177 and 202 PPC, with NNT ranging from 1012.3 and 1318.6 doses per PPC depending on when the intervention is initiated. The same strategy implemented with vaccines rather than antibiotics would target fewer than 80 PPC, with NNV exceeding 3,000 vaccines per PPC.

### Effect of reactive antibiotic prophylaxis across a range of parameters

Other parameters relating to how the strategies are implemented affect the success of the intervention (Table 4). Excluding cases that tested negative for the presence of *N. meningitidis* reduces PPC and increases NNT, but the trends in Figs 3 and 4 are unaffected, and the antibiotic prophylaxis strategy remains more efficient than the reactive vaccination strategy. See S1 File for TNT, NNT, and PPC across the full range of parameters explored.

**Table 4.** Potential effectiveness and efficiency of village-prophylaxis strategies across a range of parameters.

| Parameter | Value | TNT | PPC | NNT |
| --- | --- | --- | --- | --- |
| Baseline set | - | 181,612 | 178 | 1020 |
| Time window start | 1 | 181,612 | 198 | 917 |
|  | 7 | 181,612 | 64 | 2838 |
| Time window end | 7 | 181,612 | 128 | 1419 |
|  | 21 | 181,612 | 211 | 861 |
| Age range | 5-29 | 98,070 | 134 | 732 |
| Maximum village population | 1,000 | 55,648 | 62 | 898 |
|  | 5,000 | 163,093 | 159 | 1026 |
| Repeated doses | Yes | 287,491 | 221 | 1301 |
| Excluding cases testing negative | Yes | 181,612 | 102.6 | 1770 |

Total number treated, potentially prevented cases, and number needed to treat under a range of parameters.

## Discussion

In this outbreak in a largely rural region of Niger, there is measurable clustering of cases at the village level only after the epidemic threshold was reached, and a village-wide prophylaxis approach implemented during the epidemic targets secondary cases within villages, with a maximum of 200 out of 672 suspected cases targeted for across different parameters pertaining to implementation of the strategy.

Household prophylaxis is currently recommended in the African Meningitis Belt only outside of an epidemic. Data from this outbreak provide evidence that household prophylaxis during an epidemic can be an efficient way to target secondary cases within the household, but that such a strategy would have had minimal impact on the overall burden of disease during the outbreak. We found that clustering of cases at the household level was explained by households being in higher-burden villages, as has been observed for other infectious diseases(22). There was no evidence that household clustering was any stronger before the epidemic threshold was reached, suggesting that the strategy would target a similar number of people during an epidemic.

Previous research has focused on the effect of different epidemic thresholds on the effectiveness of reactive mass vaccination. We found that the success of the village-prophylaxis strategy is not strongly dependent on the value of the threshold used, because the threshold is used to initiate a reactive intervention. Performing surveillance at larger spatial units does not markedly improve the success of the village-wide strategy, suggesting that much of the benefit of the village-prophylaxis strategy is gained from the targeting of the villages themselves. Although including multiple villages in a round of prophylaxis can increase the number of cases targeted, the dosing of villages that would have experienced no cases leads to a general increase in NNT for these radial strategies. This seems to contrast with the finding of Maïnassara et al(17) for reactive vaccination, that health area surveillance combined with district-level vaccination was the most effective strategy; however, the difference is traceable to the difference between vaccination and antibiotic prophylaxis. Because vaccination protects individuals until the end of the season, a reactive vaccination strategy cannot get ahead of the spatial clustering in the same way.

A potential advantage of reactive prophylaxis over reactive mass vaccination is the ability to perform such a strategy within days rather than weeks of the alert threshold being reached. Similarly, the biological effect of antibiotic prophylaxis is immediate, while there is a lag between receiving a vaccination and gaining immunity. In this outbreak, prophylaxis strategies generally perform better than the equivalent reactive vaccination strategies in terms of effectiveness and efficiency because they can be triggered later and thus target more high-risk areas. The best vaccination strategy is one that targets all individuals at the beginning of the season, but such a strategy would be inefficient in a season without a large epidemic.

This study is one of a number that have assessed the clustering of meningitis cases by household in Meningitis Belt countries. Three case-control studies conducted following outbreaks reported positive or null associations between meningococcal meningitis and a household contact(23–25). In addition, several cross-sectional carriage surveys have been performed that reported the association between carriage of *N. meningitidis* and household contact of a meningococcal meningitis case(26–29). These studies generally report a positive association, although only two reached statistical significance. Finally, a longitudinal carriage study carried out during the MenAfriVac campaign found a 4.5-fold increase in acquisition rate of carriage for household contacts of a case compared to non-contacts(7).

Our finding that 4% of cases in this outbreak were secondary within a household reflects an upper bound on the proportion of infections that are household-acquired, and is similar to recent estimates for meningococcal meningitis in Western countries(30). Increased meningococcal meningitis risk to household contacts and the low proportion of meningococcal meningitis cases that are household-acquired are not inconsistent findings. Households are small and the overall population incidence rate is low, so even if the household risk ratio is high, household members’ absolute risk of meningococcal meningitis is small, and few individuals are exposed to a primary case in a household. It is thus important to understand that targeting the household is unlikely to have an impact on disease burden at the population level, even though this might be a high-risk group, when carriage prevalence and community transmission are high. In general, the effectiveness of household interventions is bounded by the proportion of infections acquired in the household, but is additionally determined by the timing of the intervention and the serial interval. A household transmission study for *N. meningitidis* carriage during an outbreak, while very challenging, would provide valuable insight into such parameters.

In analyzing this outbreak, we focused on potentially preventable cases in the absence of a comparator in which an intervention was performed, so our results have limited external comparability with other studies of meningitis outbreaks - specifically, we did not consider incomplete coverage or imperfect efficacy of prophylaxis. In addition, the effect of ciprofloxacin distribution on transmission dynamics of *N. meningitidis* is not considered, meaning that our estimates may miss some important indirect effects of administering prophylaxis on a large scale. We made a simplifying assumption that prophylaxis prevents any cases that would have occurred during a given time window, but this parameter is unknown. The focus on a single season in which an outbreak did occur limits the generalizability of our results because we did not have access to a “control” season in which there was low burden of meningococcal meningitis. Therefore, conclusions about the benefits of lower thresholds should be considered in this context.

The data on which this analysis was based consists of suspected cases reporting to health centres and hospitals in the region. As such, cases that did not present to a health centre but were still preventable are not counted in the analysis. The method for linking case data to census data was not perfect due to missing villages in the census data and villages with different names. As a result, 63 cases were excluded from the analysis due to missing or ambiguous village location and population data. Although these two effects lead to underestimation of the effect of village-wide prophylaxis, the trends observed are likely to be robust to missingness unless there is systematic bias in the presence of missingness, for example by time of year.

The recent trial of antibiotic prophylaxis in response to a meningitis epidemic showed promising results. Analysis of historical data shows that there is little household clustering of meningitis cases, and that household prophylaxis would have had limited effect on the course of the epidemic, similar to results seen in the trial. On the other hand, there is clustering of meningitis cases at the village level during an epidemic, and a reactive village-prophylaxis strategy conducted in epidemic districts can target secondary cases in villages. Our results also suggest that village-wide prophylaxis is more efficient than highly targeted reactive vaccination. However, the longer-term effectiveness of prophylaxis strategies on their own may be limited, and should thus be considered alongside reactive vaccination.

## Acknowledgments

The authors would like to thank Aimee Taylor for designing the protocol schematic in Figure 2.

## Supporting Information

**S1 Table. Clustering metrics at the household and village level, in the non-epidemic and epidemic periods.** Household and village relative risk and proportion of secondary cases, estimated in the nonepidemic and epidemic periods.

**S1 File. Supplementary results.** TNT, PPC, and NNT/NNV of reactive prophylaxis and vaccination strategies across a range of parameters.

